# Metabolic characterization of menopause: cross-sectional and longitudinal evidence

**DOI:** 10.1101/195909

**Authors:** Qin Wang, Diana L Santos Ferreira, Scott M Nelson, Naveed Sattar, Mika Ala-Korpela, Debbie A Lawlor

## Abstract

**Background:** It remains elusive whether the changes in cardiometabolic biomarkers during the menopausal transition are due to ovarian aging or chronological aging. Well-conducted longitudinal studies are required to determine this. The aim of this study was to explore the cross-sectional and longitudinal associations of reproductive status defined according to the 2012 Stages of Reproductive Aging Workshop criteria with 74 metabolic biomarkers, and establish whether any associations are independent of age related changes.

**Methods:** We determined cross-sectional associations of reproductive status with metabolic profiling in 3,312 UK midlife women. In a subgroup of 1,492 women who had repeat assessments after 2.5 years, we assessed how change in reproductive status was associated with the changes in metabolic biomarkers. Metabolic profiles were measured by high-throughput quantitative serum NMR metabolomics. In longitudinal analyses, we compared the change in metabolic biomarkers for each reproductive status category change to that in the reference of being pre-menopausal at both time points. As all women aged by a similar amount during follow-up, these analyses contribute to distinguish age related changes from those related to change in reproductive status.

**Results:** Consistent cross-sectional and longitudinal associations of menopause with a wide range of metabolic biomarkers were observed, suggesting transition to menopause induces multiple metabolic changes independent of chronological aging. The metabolic changes included increased concentrations of very small VLDL, IDL and LDL subclasses, remnant and LDL cholesterol, and reduced LDL particle size, all towards an atherogenic lipoprotein profile. Increased inflammation was suggested via an inflammatory biomarker, glycoprotein acetyls, but not via C-reactive protein. Also, levels of glutamine and albumin were increased during the transition. Most of these metabolic changes seen at the time of becoming post-menopausal remained or became slightly stronger during the post-menopausal years.

**Conclusions:** Transition to post-menopause has effects on multiple circulating metabolic biomarkers, over and above the underlying age trajectory. The adverse changes in multiple apolipoprotein-B containing lipoprotein subclasses and increased inflammation may underlie women’s increased cardiometabolic risk in post-menopausal years.

**Abbreviations:**

ALSPAC: Avon Longitudinal Study of Parents and Children
BMI: body mass index
CRP: high sensitive C-reactive protein
CVD: cardiovascular diseases
HDL: high-density lipoprotein
HRT: hormone replacement therapy
IDL: intermediate-density lipoprotein
LDL: low-density lipoprotein
SD: standard deviation
STRAW: Stages of Reproductive Aging Workshop
SWAN: The Study of Women’s health Across the Nation
VLDL: very low-density lipoprotein

## Introduction

Women in high income countries live, on average, between 35-40% of their life in the post-menopausal state, with those in low and middle-income countries also increasingly spending a high proportion of their lives post-menopausal [1]. Younger age at menopause is related to increased risk of osteoporosis, cardiovascular diseases (CVD), diabetes and premature mortality, and reduced risk of ovarian cancer and breast cancer [1-7]. Similarly, a number of disease markers, such as cardiovascular risk factors, cognitive function and bone density, have been shown to have more adverse levels in women who are post-menopausal compared with those who are pre-menopausal [8-12]. Assessment of ‘biological aging’, based on white cell DNA methylation, suggests that women experience an acceleration of biological aging with the onset of the menopause [13]. The differences in the risk markers between post- and pre-menopausal women, could thus reflect the start of biological aging and disease risk related to the change in reproductive status and its associated sex hormone changes. However, transition to post-menopause is often accompanied by the additional effects of chronological aging and midlife social adjustment [14] and in cross-sectional studies it is impossible to distinguish the effect of reproductive aging from the effect of chronological aging [11,15].

The range of diseases related to age at menopause suggest that it may have an impact on multiple metabolic pathways. Prospective studies with repeat measurements of comprehensive metabolic profiles, together with accurate characterisation of reproductive status, are required to determine this. However, to date there have been only a few prospective longitudinal studies and these have either had small sample sizes, poorly defined reproductive status, or primarily explored changes in standard lipids (total cholesterol, high-density lipoprotein (HDL) cholesterol, low-density lipoprotein (LDL) cholesterol, triglycerides), glucose, and sometimes insulin [16-19].

To study the molecular changes in response to menopausal transition and its independent effects from chronological aging, the present study investigated the cross-sectional and longitudinal associations of reproductive status defined according to the 2012 Stages of Reproductive Aging Workshop (STRAW) criteria [20] with 74 circulating metabolic biomarkers assessed primarily by high-throughput serum NMR metabolomics [21-23].

## Methods

Data from the Avon Longitudinal Study of Parents and Children (ALSPAC) were used. Full details of the recruitment, follow-up and data collection of these women have been reported previously and are available on the study website (http://www.alspac.bris.ac.uk) [24,25]. ALSPAC is a prospective population-based pregnancy cohort study that recruited 14,541 pregnancies to women resident in the South West of England between 1st April 1991 and 31st December 1992. Among them, 13,761 individual women delivered at least one live-birth and their children have been the main focus of detailed follow-up since then. Approximately 18-years after their original pregnancy a detailed assessment of mothers was completed [25], and the current study is based on the 3,312 mothers who attended a clinic assessment between December 2008 and June 2011 (median age (IQR): 48 (45, 51)) and a subgroup of those women (N = 1,492) who attend a second follow-up assessment approximately 2.5 years later (median age (IQR): 51 (48, 54)). Figure 1A illustrates the flow of women into eligible and analyses groups. Baseline characteristics of participating women are shown in Table 1.

**Table 1.** Baseline characteristics for ALSPAC mums.

| **Characteristics** | **Pre-menopausal**<br>**(N = 2031)** | **Menopausal transition**<br>**(N = 620)** | **Post-menopausal**<br>**(N = 661)** |
| --- | --- | --- | --- |
| Age (years) | 46 [44-48] | 50 [48-52] | 54 [51-56] |
| BMI (kg/m<sup>2</sup>) | 25 [23-29] | 25 [22-29] | 25 [22-29] |
| Height (cm) | 164 [160-168] | 164 [160-168] | 163 [160-167] |
| Fat mass (kg) | 25 [19-32] | 24 [18-32] | 25 [19-32] |
| Lean mass (kg) | 41 [38-44] | 41 [38-44] | 40 [37-43] |
| Trunk fat mass (kg) | 13 [ 9-18] | 12 [ 9-17] | 13 [ 9-17] |
| Systolic blood pressure (mmHg) | 115 [109-124] | 116 [109-125] | 117 [110-127] |
| Diastolic blood pressure (mmHg) | 70 [66-76] | 71 [66-77] | 72 [66-78] |
| Educated to university level n/N (%)<sup>a</sup> | 301/1887 (16) | 149/574 (26) | 159/611 (26) |
| White European n/N (%)<sup>a</sup> | 1835/1881 (98) | 563/573 (98) | 588/607 (97) |
| Lipid-lowering medication n/N (%) | 29/2031 (1) | 9/620 (1) | 13/661 (2) |

Values are presented as median [interquartile range] unless otherwise stated.
<sup>a</sup> n/N = number with the characteristic divided by the total number in the group with no missing data on that variable.

**Figure 1.**
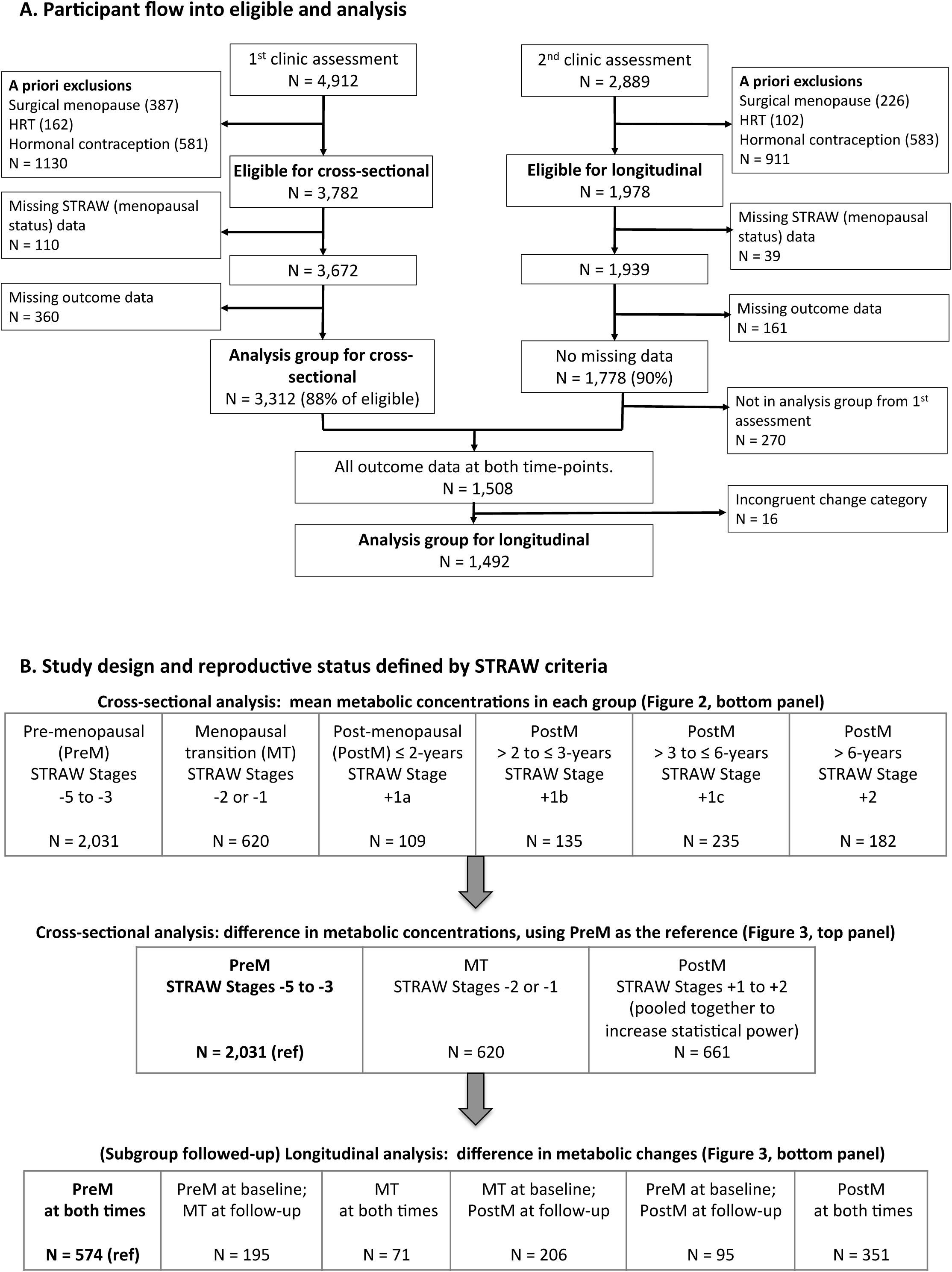
Participant flow and study design.

### Assessment of reproductive status (exposure)

At both clinic assessments women were asked a range of questions concerned with their menstrual cycle including its frequency and regularity and date of last menstrual period, which enabled them to be categorised according to the 2012 Stages of Reproductive Age Workshop STRAW criteria [20]. There are ten STRAW stages, which are grouped into three larger groups: (i) Reproductive (referred to as pre-menopause in this paper), which includes categories -5 (beginning at menarche and characterised by variable to regular menstrual cycles), -4 (regular), -3b (regular) and -3a (start of subtle changes in cycle length); (ii) menopausal transition, which includes categories -2 (variable length of cycle) and -1 (categorised by episodes of amenorrhoea of ≥ 60 days); (iii) Post-menopause, which begins at the last menstrual period and includes +1a (up to the first two years since the last menstrual period), +1b (>2 to ≤3 years), +1c (>3 to ≤6 years), and 2 (> 6 years). To increase statistical power, in all analyses we combined women in all four pre-menopausal categories (i.e. -5 to -3) into one pre-menopausal category and combined those in the two menopausal transition categories (-2 and -1) into one menopausal transition category (Figure 1B). Women were also asked about previous hysterectomy, oophorectomy, endometrial ablation or radio- or chemotherapy related to reproductive organs (together defined as surgical menopause), and about the use of hormone replacement and contraception. We excluded *a priori* those women who had experienced a surgical menopause, and those using hormone replacement or hormonal contraception, at baseline (for both analyses) and also at follow-up (for longitudinal analyses), in order to assess naturally occurring changes across reproductive categories (Figure 1A). The remaining women, who had completed information on STRAW data and metabolic profiles, were then classified into one of the six mutually exclusive reproductive status groups based on the STRAW criteria (Figure 1B, **top row**). None of the women were pregnant at baseline or follow-up.

### Assessment of metabolic profiling, anthropometry, and blood pressure (outcomes)

At both assessments blood samples were taken after an overnight fast for those examined in the morning and a minimum 6-hours for those seen after 14.00. Blood samples were immediately spun and frozen at –80°C and all of the assays completed for this study were undertaken within 3-years of storage and with no previous freeze-thaw cycles. A high-throughput NMR metabolomics platform was used to quantify 73 serum lipid and metabolite measures. The platform applies a single experimental setup, which allows for the simultaneous quantification of routine lipids, 14 lipoprotein subclasses and individual lipids transported by these particles, multiple fatty acids, glucose and various glycolysis precursors, ketone bodies and amino acids in absolute concentration units. Details of this platform have been published previously [21,22], and it has been widely applied in genetic and observational epidemiological studies [23,26-34]. High sensitive C-reactive protein (CRP) was measured by automated particle-enhanced immunoturbidimetric assay (Roche UK). Together, these 74 metabolic measures are defined as the primary outcomes and constitute the circulating metabolic profile.

In addition, we also analysed the associations of reproductive status with anthropometric measures and blood pressure. Weight and height (used to calculated the body mass index (BMI)) were measured in light clothing and unshod. Weight was measured to the nearest 0.1kg using Tanita scales; height was measured to the nearest 0.1cm using a Harpenden stadiometer. A lunar prodigy DXA scan was used to measure total body fat-, trunk fat- and total body lean-mass. Seated blood pressure was measured with the woman at rest, her arm supported and the correct cuff-size (after measurement of arm circumference), using an Omron M6 upper arm monitor. Two measurements were undertaken and their mean used.

### Assessment of potential confounders

Age was recorded at each assessment. Educational attainment and ethnicity, which were considered potential confounders [17], were obtained by questionnaire when the women were originally recruited [25]. Use of lipid-lowering medication, which affects lipid concentrations [28], was assessed by questionnaire. Fat mass, which affects the systemic metabolic profile [33], was also considered as a potential confounder.

#### Statistical analyses

The metabolic measures were scaled to standard deviation (SD) units (by subtracting the mean and dividing the standard deviation of all women included in the baseline analysis). This scaling allows easy comparison of multiple metabolic measures with different units or with large differences in their concentration distributions.

In cross-sectional analyses, the heatmap of metabolic profile across the age groups was compared to the heatmap of metabolic profile across the reproductive status groups. Baseline women were categorized into different age groups, largely defined in 1-year age categories (at the extremes of age range groups were collapsed because of small numbers: 34- to 38-years and 59- to 63-years). Similarly, the same women were categorized into six reproductive status groups according to STRAW criteria (Figure 1B, **top row**). Heatmap colours illustrate the mean concentration of metabolic outcomes in each of these groups. Then, linear regression models were used to quantify the differences in the metabolic concentrations across the reproductive status groups, using pre-menopausal women as the reference group. To increase statistical power, all the four post-menopausal groups were pooled together as a single post-menopausal group (Figure 1B, **middle row**).

In longitudinal analyses, a subgroup of women, who had repeated exposure and outcome measures 2.5 years later, were categorized into six groups based on their baseline and follow-up reproductive status (Figure 1B, **bottom row**). Firstly, the mean difference between baseline and follow-up for each outcome in all six categories was calculated. Then, linear regression models were used to estimate the differences in mean differences between five change categories compared with those who were pre-menopausal at both baseline and 2.5 years follow-up (the reference group which primarily reflects age related changes). Under the assumption that changing from one reproductive status category to another is not associated with other changes that affect outcomes (beyond the confounders that we adjust for), this ‘difference in differences’ is a valid means of testing causal effects with longitudinal observational data [35]. As the metabolic changes in the reference groups estimates the metabolic changes primarily due to 2.5-y chronological aging, the results of ‘difference in differences’ can be interpreted as the extent to which the longitudinal changes are deviated from the underlying age trajectory. For example, null associations illustrate the metabolic changes occurred during the follow-up were likely to be only due to age-related change (and little or no impact from changing reproductive status), while pronounced associations demonstrate that the reproductive status changes were over and above the age effect.

Due to the correlated nature of the metabolic biomarkers, over 95% of the variation in the 74 metabolic biomarkers was explained by 19 principal components. Therefore, multiple testing correction, accounting for 19 independent tests using the Bonferroni method, resulted in P < 0.002 (0.05/19) being denoted as statistically significant [32,34]. All analyses were undertaken in the statistical software package R (version 3.4.0).

### Additional analyses

In the main analyses, cross-sectional and longitudinal associations (linear regression models) were adjusted for baseline age. In additional analyses, cross sectional analyses were further adjusted for education, ethnicity, lipid-lowering medication, fat mass and height, and longitudinal analyses were additionally adjusted for baseline education, ethnicity and 2.5-year change in lipid-lowering medication, fat mass and height. Cross-sectional and longitudinal analyses with anthropometric and blood pressure outcomes were also undertaken. Whilst our focus here is on metabolic profiles these analyses allowed comparisons with previous studies of the association of reproductive status with established CVD risk factors. Lastly, as both natural menopause and surgical menopause indicate the decline of ovarian function, we examined, in those women who had been previously excluded, the cross-sectional and longitudinal associations of surgical menopause with metabolic profiles to explore whether surgical menopause displayed similar association-pattern to those seen for natural menopause.

## Results

Of the 3,782 eligible women attending the baseline assessment (i.e. after *a priori* exclusions) 3,312 (88%) had complete data on reproductive status and systemic metabolic measures; equivalent numbers for the follow-up assessment were 1,978 and 1,778 (90%) (Figure 1A). A total of 1,508 women had complete data at both time points. The 3,312 women with complete baseline data were included in cross-sectional analyses and 1,492 with complete baseline and follow-up data, and who had valid change in reproductive status over time (16 women had menopause status change that was not plausible, e.g. appearing to change from post-menopause to menopause transition), were included in longitudinal analyses. Broadly similar baseline characteristics were observed for those women included in the analyses, women excluded *a priori* and those excluded because of missing data (**Additional file 1: Table S1**).

### Metabolic heatmap of chronological aging and menopause

Figure 2 **(top panel)** shows mean metabolic concentrations for women aged from late 30s to early 60s. Concentration of multiple lipoprotein particles, including intermediate-density lipoprotein (IDL) and LDL subclasses, and the individual lipids (triglycerides, cholesterol and phospholipids) transported by these particles, increased with chronological age. The patterns of associations with these measures were similar, with the metabolic concentrations on average being the lowest during late 30s to mid 40s, then slightly increased during mid 40s to early 50s, and became substantially higher from early 50s. Similar chronological aging patterns were observed for apolipoprotein A1 and B, total cholesterol and phospholipids, absolute fatty acid concentrations, the ratio of omega-3 fatty acids and its subclass DHA, as well as multiple non-lipid measures including glutamine, tyrosine, glucose, and citrate. Given the average menopausal age was at 51 in European ancestry [36], it is plausible that the marked increase around age 50 in these metabolic profiles were associated with reproductive status change. Weaker increases around and after the age of menopause were seen for very low-density lipoprotein (VLDL) and HDL subclasses (except for very small VLDL), the individual lipids in these particles, and total triglycerides. LDL particle sizes became smaller with increasing age. Over and above the pattern observed around the menopausal age, we also observed an increase and then decrease in most VLDL subclasses, total and VLDL triglycerides, all branch-chain amino acids (and some of the other amino acids), lactate, pyruvate, albumin, glycoprotein acetyls and CRP roughly around ages 39 to 43. The biological reasons or confounding factors, which could explain the increase in these metabolic measures between these ages, are however unclear and we have insufficient data on women in their earlier 30’s to fully describe the nature of these changes.

**Figure 2.**
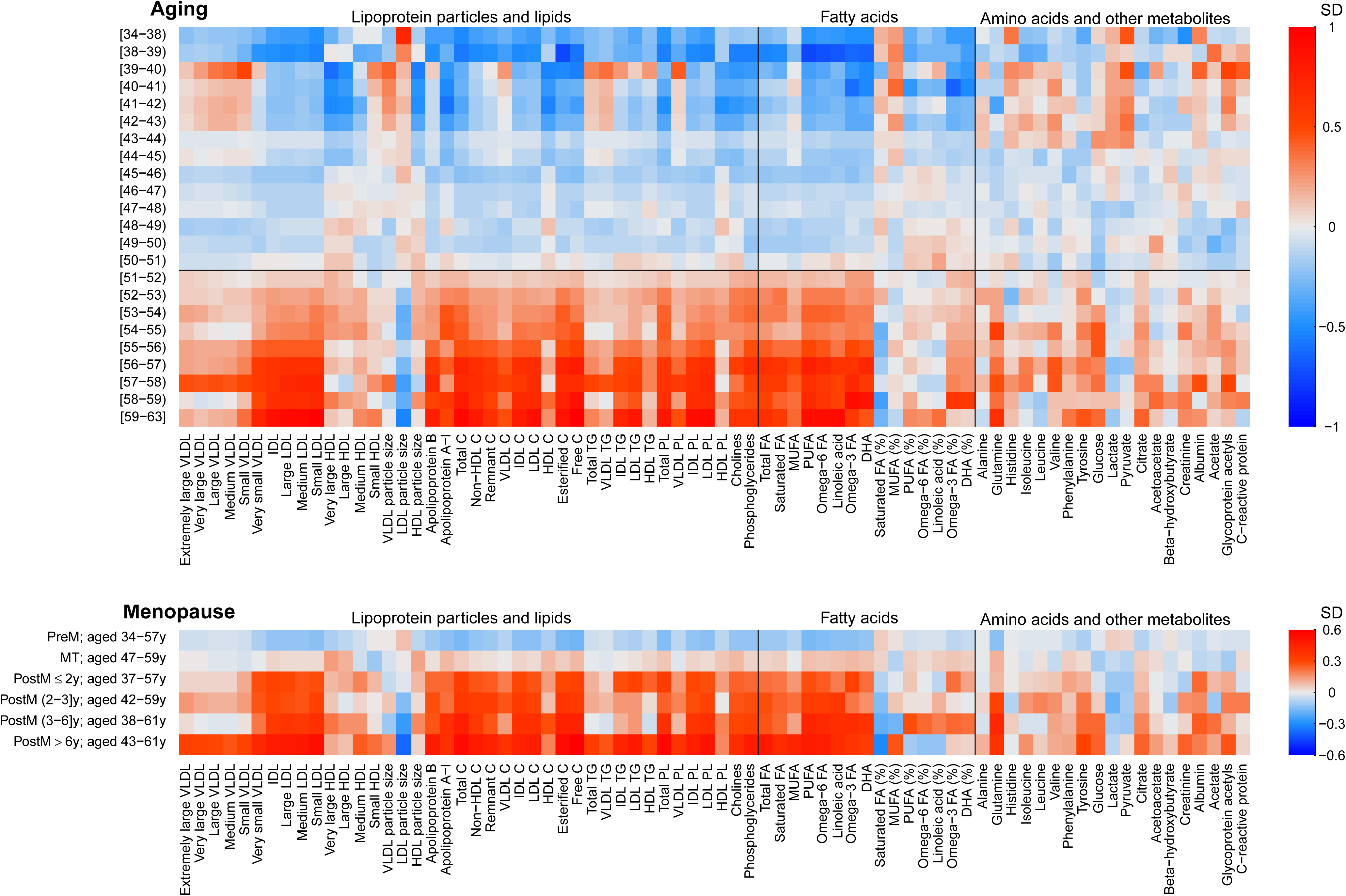
Heatmap of metabolic concentrations across age groups (top panel) and across reproductive categories (bottom panel). Top panel: age groups are largely in 1-year age categories (at the extremes of age range, groups were collapsed because of small numbers: 34- to 38-years and 59- to 63-years). For each metabolic measure, mean metabolite concentrations across each age group are displayed in colours. The horizontal black line marks the typical age of natural menopause in Europe. **Bottom panel:** reproductive categories are defined according to the STRAW criteria. For each metabolic measure, mean metabolite concentrations across each menopausal category are displayed in colours. The age range of each reproductive category is shown on the y-axis. As the metabolic measures were scaled to SD units, both analyses (ageing and menopause) thus used the population mean of all women at baseline as the reference value, marked as white boxes in the plots. The metabolic measures in both panels are plotted in the same order. C, cholesterol; TG, triglycerides; PL, phospholipids; FA, fatty acids; MUFA, monounsaturated fatty acids; PUFA, polyunsaturated fatty acids; DHA, docosahexaenoic acid.

Figure 2 **(bottom panel)** illustrates the mean metabolic concentrations across the reproductive groups. Multiple metabolic concentrations were substantially greater in post-menopausal compared with pre-menopausal women, with a similar pattern to that seen for the marked increase in metabolites at early 50’s. Given the reproductive groups are based on individual menstrual cycle patterns and each group has within it a large variation in age (and overlap across categories), the similar metabolic patterns suggest that the metabolic aberrations are associated with change in reproductive status over and above chronological aging.

### Cross-sectional associations of reproductive status

Figure 3 **(top panel)** shows the cross-sectional, age-adjusted, associations of reproductive status with 74 metabolic measures. In age adjusted analyses, women who were post-menopausal, compared with those who were pre-menopausal had higher concentration of multiple lipoprotein lipids, particularly those IDL and LDL related measures, increased concentrations of fatty acids, and several non-lipid measures. In contrast, there were only minor metabolic differences between women in the menopausal transition and those who were pre-menopausal.

**Figure 3.**
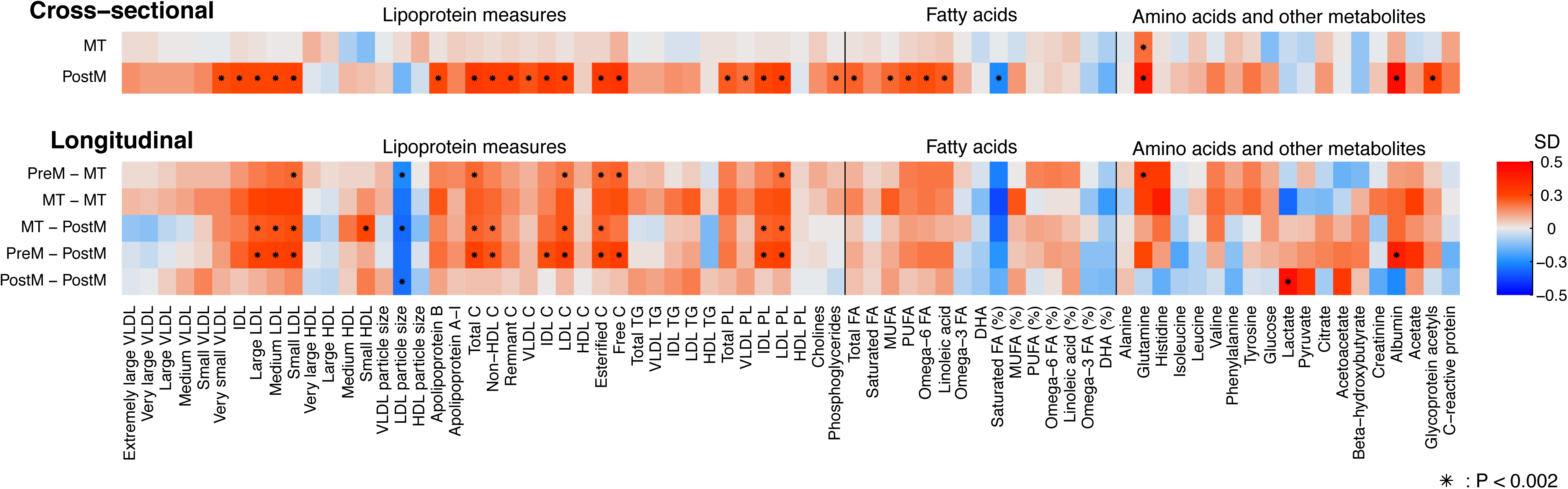
Cross-sectional (top panel) and longitudinal (bottom panel) associations of reproductive categories with 74 metabolic measures. Top panel. In cross-sectional analysis, women were categorised into pre-menopausal, menopausal transition and post-menopausal groups. Using pre-menopausal women as the reference group, the colours define the SD-differences in each metabolite concentration between the other two groups and the reference group (Figure 1B). **Bottom panel.** In longitudinal analysis, women were categorized into 6 groups based on their baseline and follow-up reproductive status. Using those women who were pre-menopausal at both time-points as the reference group, the colours illustrate the SD-differences in the concentration change during the follow-up between the other five groups and the reference group (Figure 1B). Cross-sectional and longitudinal associations were adjusted for baseline age. The metabolites are plotted in the same order as in Figure 2. C, cholesterol; TG, triglycerides; PL, phospholipids; FA, fatty acids; MUFA, monounsaturated fatty acids; PUFA, polyunsaturated fatty acids; DHA, docosahexaenoic acid.

### Longitudinal changes in response to reproductive status change

Figure 3 **(bottom panel)** shows the longitudinal differences in mean differences for each reproductive status change category compared with the reference category of being pre-menopausal at both time points, reflecting the amount by which outcomes changed over the follow-up period in response to change in the reproductive status. Within each category of reproductive status ‘change’, including the reference category of women who remained pre-menopausal at both time points, metabolite concentrations changed as all women aged 2.5 years during the follow-up (**Additional file 1: Figure S1**). In comparison with those women who were pre-menopausal at both time-points, greater increases in IDL and LDL-related measures, larger decrease in LDL particle size and greater changes in multiple fatty acids and non-lipid measures were consistently observed for the four groups of women who changed from pre- to menopausal transition, from menopausal transition to a 2.5-y greater exposure to menopausal transition, from menopausal transition to post-menopausal and from pre- to post-menopausal. These results suggest metabolic changes, over and above the underlying age trajectory, occurring during the transition period from pre- to post-menopause. By contrast, the metabolic changes for those women who were post-menopausal at both time-points were less notably different to those who were pre-menopausal at both time-points.

### Summary of cross-sectional and longitudinal associations of change in reproductive status

Figure 4 **and Table S2 (Additional file 1)** summarize and compare the cross-sectional and longitudinal associations of reproductive status and its change with metabolic traits. The cross-sectional results are differences in mean concentrations comparing post- to pre-menopausal women. The longitudinal association results were meta-analyzed across four groups of women – those changing from pre- to menopausal transition, menopausal transition to 2.5 years later still in menopausal transition, menopausal transition to post-menopause, and pre- to post-menopause – compared with those who were pre-menopausal at both time point. The fixed-effect meta-analysis was done to increase statistical power, given that these four groups displayed broadly similar metabolic association patterns (Figure 3 **lower panel**) and that all of them represent, at least partly, the transition process from pre- to post-menopause. A meta-analysis was undertaken as it would not have been possible to combine these women in to one single change group given they changed through different categories.

Longitudinal associations of menopause were broadly similar as the cross-sectional associations (Figure 4). Consistent associations were observed for increased concentrations of very small VLDL, IDL and LDL particles, as well as the cholesterol and phospholipids transported in these particles, whereas there was little robust evidence from cross-sectional or longitudinal analyses in supporting the associations with triglycerides. Large increases in total cholesterol, remnant cholesterol, total phospholipids, apolipoprotein B, and apolipoprotein A-1 were also observed. However, there were only weak associations with HDL-related measures. While LDL particle size decreased as women changed from pre- to post-menopause, weak changes were seen for VLDL and HDL particle sizes. Total fatty acids, omega-6 fatty acids and its subclass of linoleic acid were increased, and the ratio of saturated fatty acids relative to total fatty acids decreased as women transitioned from pre- to post-menopause. For the non-lipid measures, large increases were observed for glutamine, histidine, valine, albumin and acetate as women became post-menopausal. Interestingly, glycoprotein acetyls (an inflammatory marker) increased with transition from pre- to post-menopause, whereas change in reproductive status was not robustly associated with the other inflammatory marker C-reactive protein in either longitudinal or cross-sectional analyses (the 95% confidence interval included the null in the longitudinal analyses). The magnitudes of change in these metabolic concentrations as women became post-menopausal were broadly similar (∼ 0.2 SDs).

**Figure 4.**
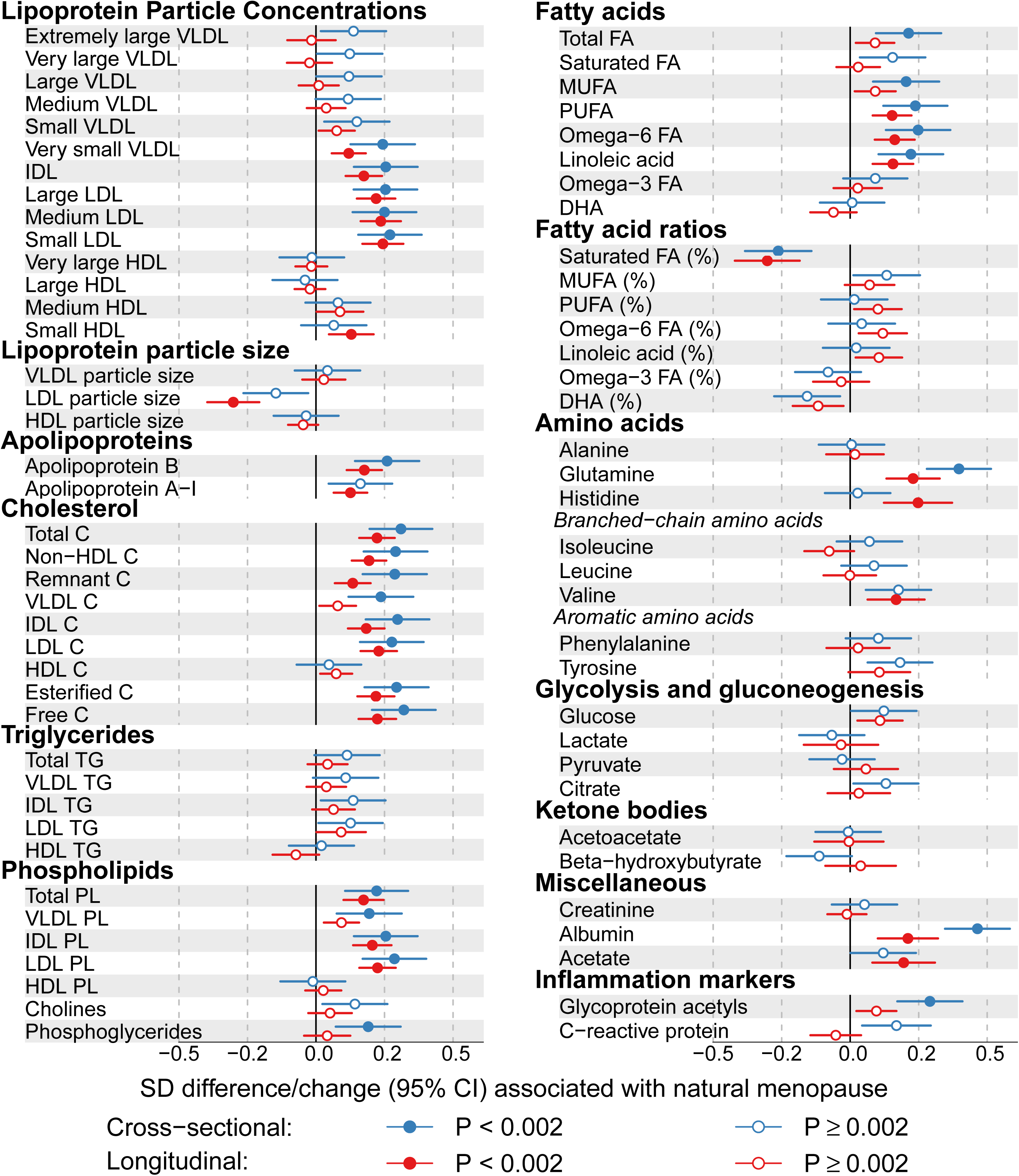
Cross-sectional (blue) and longitudinal (red) associations of natural menopause with 74 metabolic measures. The cross-sectional associations are differences in mean metabolites comparing post-menopausal to pre-menopausal women. The longitudinal associations are the differences in mean differences over time comparing four groups meta-analyzed together (Pre-menopause to Menopause Transition (PreM – MT), Menopause Transition at both times (MT – MT), Menopause Transition to Post-Menopause (MT – PostM), Pre-Menopause to Post-Menopause (PreM – PostM)) to Pre-menopause at both times (preM-preM; the reference group). The meta-analysis of the four groups, each of which at least partly represent the transition process from pre-menopausal to post-menopausal, was used to increase statistical power. Cross-sectional and longitudinal associations were adjusted for baseline age. C, cholesterol; TG, triglycerides; PL, phospholipids; FA, fatty acids; MUFA, monounsaturated fatty acids; PUFA, polyunsaturated fatty acids; DHA, docosahexaenoic acid.

### Additional analyses

Cross-sectional associations of reproductive status with circulating metabolic measures remained largely unchanged when further adjusted for ethnicity, education, fat mass, height and lipid-lowering medication (**Additional file 1: Figure S2**). Longitudinal associations also remained similar when adjusted for additional potential confounders, including baseline ethnicity and education and 2.5-y change in fat mass, height and lipid-lowering medication (**Additional file 1: Figure S2**). Additional analyses of reproductive status with anthropometry and blood pressures were also conducted (**Additional file 1: Figure S3**). In cross-sectional analyses height and lean mass were lower, and total and truncal fat mass, and diastolic blood pressure higher in post-compared to pre-menopausal women. However, with the exception of the inverse association with lean mass, these differences did not reach our multiple testing p-value threshold and were not replicated in longitudinal analyses. Further, there was no strong evidence for association of reproductive status change with BMI, waist circumference or systolic blood pressure in either cross-sectional or longitudinal analyses. Associations of surgical menopause with multiple metabolic measures were seen in cross-sectional analysis, whereas longitudinal results were imprecisely estimated and difficult to reliably interpret due to the small number of women who underwent a surgical menopause during the follow-up (**Additional file 1: Figure S4**).

## Discussion

In this study of 3,312 midlife women, cross-sectional and longitudinal analyses were used to determine the relation of reproductive status with comprehensive metabolic profiling. Cross-sectional analyses showed that being post-menopausal was associated with a wide range of circulating metabolic measures, including multiple established and emerging biomarkers for type 2 diabetes, cardiovascular diseases and all-cause mortality [26,27,37]. The ‘difference in mean differences’ in the longitudinal analyses displayed broadly consistent association-patterns with the cross-sectional analysis, suggesting the observed metabolic aberrations were due to the effect of reproductive status change, over and above age-related changes. These metabolic changes were primarily towards an atherogenic lipid profile, with increased concentrations of small VLDL, IDL and LDL subclasses, higher levels of all cholesterol measures except HDL cholesterol, increased apolipoprotein-B and decreased LDL particle size. We also found that becoming post-menopausal resulted in decreased proportion of saturated fatty acids, higher concentrations of glutamine, valine, albumin, acetate and glycoprotein acetyls. The majority of these metabolic changes appear to persist, or possibly increase slightly, over time after becoming post-menopausal.

One previous study examined cross-sectional associations of age with similar metabolic profiles to those assessed here and showed that around the average age at menopause for European women metabolic profiles changed more so in women than men [8], with patterns that are broadly consistent with the findings we find with detailed characterisation of reproductive status and longitudinal analyses. The Study of Women’s health Across the Nation (SWAN) is the previous largest study (N = 1,054) to examine prospective associations of reproductive status change with changes in established risk factors [17]. In consistent with our study, they found longitudinal increases in total cholesterol, LDL cholesterol and apolipoprotein-B on becoming post-menopausal, that were independent of chronological aging. As in our study, SWAN also found that changes in BMI, blood pressure, triglycerides, HDL cholesterol and C-reactive protein in women’s midlife were small and/or largely expected to be related to age, rather than change in reproductive status. Although there is little evidence in supporting the change in C-reactive protein in both SWAN and our longitudinal analysis, our results suggested that the change from pre- to post-menopause increases the concentration of glycoprotein acetyls, a biomarker for low-grade chronic inflammation, which has been shown to be positively associated with diabetes, cardiovascular disease, and premature mortality [26,38-41]. Thus, becoming post-menopausal may result in increased low-grade inflammation over and above age effects. The overall consistent findings between our study and the SWAN study (which used a multilevel model analytical approach with a greater number of repeat measures) for the established risk markers also suggest that the ‘difference in mean differences’ approach taken here is a valid way to assess the metabolic consequences of change in reproductive status that is independent of chronological aging.

Besides the established lipid and inflammatory markers, our results provide longitudinal evidence for a potential impact of becoming post-menopausal on lipoprotein particles, fatty acids, amino acids and other metabolites. As women changed from pre- to post-menopause, the atherogenic lipoprotein particles, including the remnant (small VLDL + IDL) and LDL particle concentrations were increased, together with decreased LDL particle size. These changes are likely to predispose post-menopausal women towards higher CVD risk. Previous studies have reported that branched-chain and aromatic amino acids are predictive of type 2 diabetes, and that their circulating levels increase in response to weight gain and increases in insulin resistance [33,42]. Transition to post-menopause had only a weak effect on these amino acids, except for valine. These findings suggest that the adverse metabolic changes observed here are unlikely to be entirely mediated through weight gain or increased insulin resistance during the transition period, a finding consistent with our observation of little robust evidence for a relation of change in reproductive status with change in adiposity measurements, or change in the longitudinal effects on metabolic traits with adjustment for both baseline and follow-up fat mass and glucose. Interestingly, transition to post-menopause was associated with decreased proportion of saturated fatty acids and increased concentration of glutamine and albumin, all of which are associated with lower cardiometabolic risk [26,37,43]. However, the casual role of these emerging biomarkers for disease risks remains unclear.

Menopause is associated with a decrease in estradiol, beginning during the transition phase (∼ 2 years before a woman’s final menstrual period), with levels plateauing at a low value by ∼ 2 years post-menopause, and a mirror pattern of increasing FSH over the same period [44].The metabolic changes observed here may thus reflect sex and gonadotropin hormonal changes relating to the menopausal transition. Previous studies have shown exogenous estrogen alone, or in combination with progestogens (as hormone replacement therapy (HRT) or combined hormonal contraception), is associated with variation in lipid levels [30,45]. Of relevance to our findings, post-menopausal women using HRT were found to have lower LDL-cholesterol in comparison with those not using HRT [45]. However, these associations with exogenous hormones may not be causal, and it remains unclear to what extent HRT will associate with non-lipid biomarkers, e.g. fatty acids and amino acids. Several studies have recently found exogenous hormones [30] and also reproductive status change, including pregnancy [31] and menopause (as studied here) to be associated with changes in a wide-range of circulating metabolites. Further research to understand the relationship of circulating levels of reproductive hormones at different stages of the lifecourse with changes in comprehensive metabolic profiles is important to better understand the extent to which sex hormones influence metabolism during different reproductive stages of life.

Overall, consistent and prominent cross-sectional and longitudinal metabolic associations were seen with transition from pre- to post-menopause, whilst weaker associations between pre-menopause and menopause transition were noticed for cross-sectional associations in comparison with longitudinal associations. Previous prospective studies have suggested that metabolic effects of the changing from the pre-menopause to menopausal transition stage, occur mostly as a result of changing to the late stage of the menopause transition, with only weak/null metabolic effects with the change from pre-menopause to the early phase of menopausal transition [16,17]. Thus, whilst small numbers meant having to combine women who were in the early and late menopausal transition categories, this may have limited our ability to detect any specific metabolic differences between late menopausal transition and pre-menopausal women in the case of cross-sectional analysis.

The key strengths of this study are its detailed information on reproductive status and comprehensive metabolic profiles collected on two occasions prospectively in large numbers. Our results are broadly supported by the previous large-scale cross-sectional and longitudinal studies of established risk factors [8,17], however we acknowledge that replication of the associations with the emerging metabolic risk factors that we have assessed here would be important. Our results suggest that majority of the metabolic changes seen at the time of becoming post-menopausal persist over time, and for some may continue to change at a slow pace. However, the mean age of women at baseline and follow-up were 48 and 51 years, respectively, and further repeat assessments in these women would be valuable to verify this, and to determine how long such changes may continue. These women are currently too young to have experienced cardiovascular disease events, osteoporotic fractures or cancers in sufficient numbers for us to be able to examine the extent to which the menopause related changes that we have observed in metabolic profiles relate to future disease outcomes. With continued follow-up of these women that would be possible. The women in this study are largely of European origin and we cannot assume that findings generalise to other groups.

In conclusion, when women become post-menopausal, they experience changes in relation to lipoprotein metabolism, fatty acids, amino acids and inflammation. These metabolic changes are age-independent and potentially underlie the relationship between menopause and cardiometabolic diseases. Detailed understanding of the molecular changes occurred during menopausal transition may lead to lifestyle or therapeutic opportunities to diminish the adverse metabolic effects on women during their post-menopausal life.

### Declarations

#### Ethics approval and consent to participate

Ethical approval was obtained from the ALSPAC Law and Ethics Committee and the UK National Health Service Local Research Ethics Committee (references: 08/H0106/96 and 11/SW/0110). Women provided informed written consent for their assessments and use of the blood samples.

#### Consent for publication

Not applicable.

#### Availability of data and material

All ALSPAC data are available to scientists on request to the ALSPAC Executive via this website, which also provides full details and distributions of the ALSPAC study variables: http://www.bristol.ac.uk/alspac/researchers/access/. Consistent with other studies funded by UK funders ALSPAC uses a business model to offset the expense of preparing and supporting data access. The ALSPAC data management plan (available here: http://www.bristol.ac.uk/alspac/researchers/data-access/documents/alspac-data-management-plan.pdf) describes in detail the policy regarding data sharing, including the real costs for providing data. The system for accessing data via the executive applies to all researchers including ALSPAC investigators and scientists at the University of Bristol. The executive do not scrutinise the proposed science by those wishing to access data, nor do they check for scientific overlap with other data requests (all data requests are published online - https://proposals.epi.bristol.ac.uk/); requests are only refused if the necessary data are not available or there are concerns that the proposed research might bring the study into disrepute. Restrictions (including collapsing categories) might be applied to some variables if cell numbers are small. DAL was a member of the ALSPAC executive from April 2007 to June 2017. An independent scientific advisory board reviews the very small number of data access requests that are declined. The study website contains details of all the data that is available through a fully searchable data dictionary (http://www.bris.ac.uk/alspac/researchers/data-access/data-dictionary/).

#### Author’s contribution

All listed authors meet the requirements for authorship. QW, MAK and DAL designed this study and agreed the analysis plan. QW undertook analyses. DSLF provided support with data management and SNM and NS supervised the laboratory assays for CRP. QW, MAK and DAL wrote the original draft of the study, with all other authors providing comments. DAL acts as guarantor.

## Acknowledgements

We are extremely grateful to all the families who took part in this study, the midwives for their help in recruiting them, and the whole ALSPAC team, which includes interviewers, computer and laboratory technicians, clerical workers, research scientists, volunteers, managers, receptionists and nurses.

## Additional files

Additional file 1: Supplementary tables and figures in support our results and findings. (PDF)

